# Highly functional virus-specific cellular immune response in asymptomatic SARS-CoV-2 infection

**DOI:** 10.1101/2020.11.25.399139

**Authors:** Nina Le Bert, Hannah E Clapham, Anthony T Tan, Wan Ni Chia, Christine YL Tham, Jane M Lim, Kamini Kunasegaran, Linda Tan, Charles-Antoine Dutertre, Nivedita Shankar, Joey ME Lim, Louisa Jin Sun, Marina Zahari, Zaw Myo Tun, Vishakha Kumar, Beng Lee Lim, Siew Hoon Lim, Adeline Chia, Yee-Joo Tan, Paul Anantharajah Tambyah, Shirin Kalimuddin, David Lye, Jenny GH Low, Lin-Fa Wang, Wei Yee Wan, Li Yang Hsu, Antonio Bertoletti, Clarence C Tam

**Author notes:** These authors contributed equally (Antonio Bertoletti, Clarence C Tam). One Sentence Summary: Virus-specific T cells secrete high levels of IFN-γ and IL-2 in asymptomatic SARS-CoV-2 infection.

## Abstract

The efficacy of virus-specific T cells in clearing pathogens involves a fine balance between their antiviral and inflammatory features. SARS-CoV-2-specific T cells in individuals who clear SARS-CoV-2 infection without symptoms or disease could reveal non-pathological yet protective characteristics. We therefore compared the quantity and function of SARS-CoV-2-specific T cells in a cohort of asymptomatic individuals (n=85) with that of symptomatic COVID-19 patients (n=76), at different time points after antibody seroconversion. We quantified T cells reactive to structural proteins (M, NP and Spike) using ELISpot assays, and measured the magnitude of cytokine secretion (IL-2, IFN-γ, IL-4, IL-6, IL-1β, TNF-α and IL-10) in whole blood following T cell activation with SARS-CoV-2 peptide pools as a functional readout. Frequencies of T cells specific for the different SARS-CoV-2 proteins in the early phases of recovery were similar between asymptomatic and symptomatic individuals. However, we detected an increased IFN-γ and IL-2 production in asymptomatic compared to symptomatic individuals after activation of SARS-CoV-2-specific T cells in blood. This was associated with a proportional secretion of IL-10 and pro-inflammatory cytokines (IL-6, TNF-α and IL-1β) only in asymptomatic infection, while a disproportionate secretion of inflammatory cytokines was triggered by SARS-CoV-2-specific T cell activation in symptomatic individuals. Thus, asymptomatic SARS-CoV-2 infected individuals are not characterized by a weak antiviral immunity; on the contrary, they mount a robust and highly functional virus-specific cellular immune response. Their ability to induce a proportionate production of IL-10 might help to reduce inflammatory events during viral clearance.

## Introduction

Characterization of adaptive immunity mounted against SARS-CoV-2 is crucial for understanding its role in protection or pathogenesis. Antibodies and T cells act together to reduce the spread of virus within the host and to eradicate the pathogen from infected cells. However, the protective immune response can also trigger pathological processes characterized by localized or systemic inflammatory events. Inflammation and tissue damage can result from the direct lysis of infected cells by virus-specific antibodies and T cells, or from the release of inflammatory mediators produced by the infected cells and activated myeloid cells. These scenarios have been reported in the pathogenesis of COVID-19 *(1)*. In more severe cases, systemic high levels of inflammatory cytokines (IL-1β, IL-6), presence of activated monocytes in the circulatory compartment *(2, 3)* and in the lung *(4)* co-exist with virus-specific antibodies and T cells *(5)*. Thus, the question of whether virus-specific antibodies or T cells are preferentially mediating protection or damage remains open. Antibodies against SARS-CoV-2 Spike (S) protein have protective ability in vitro, but their titers in COVID-19 patients have been reported to be positively correlated with disease severity *(6, 7)*. Similarly, a direct relation with disease severity has been reported in studies of SARS-CoV-2-specific T cell frequencies in COVID-19 patients. A broader and quantitatively more robust SARS-CoV-2-specific T cell response has been demonstrated in convalescent cases of severe COVID-19 in comparison to mild cases *(8)*. However, the positive relation between SARS-CoV-2-specific T cell quantity and disease severity was not confirmed in recent studies measuring SARS-CoV-2-specific T cells in the early phases of COVID-19. Early induction and more robust SARS-CoV-2-specific CD4 and CD8 T cell response has been associated with milder disease development *(9, 10)*. In addition, a protective role of T cells has been demonstrated in animal models of Coronavirus infections, in which virus-specific T cells clear the virus with limited lung pathology *(11, 12)*.

In this work we expand our investigation into virus-specific T cells in SARS-CoV-2 infection by studying asymptomatic SARS-CoV-2 infected individuals. They constitute a large proportion of infected individuals *(13)* and should hold the key to understanding the immune response capable of controlling the virus without triggering pathological processes. However, current knowledge of their antiviral immunity is limited. SARS-CoV-2-specific antibodies *(14)* and T cells are induced in asymptomatic individuals *(15)*, but the reported lower magnitude of antibodies *(14)* and T cell responses *(15, 16)* has been interpreted as a sign that asymptomatic individuals are mounting a normal innate *(17)*, but a weak adaptive antiviral immunity *(14)*. Since antibody titers and frequency of SARS-CoV-2-specific T cells in the blood can undergo dynamic changes after viral clearance *(10)*, we reasoned that a proper quantitative comparison between SARS-CoV-2-specific T cells of symptomatic and asymptomatic individuals should be performed at similar timepoints post infection. The absence of symptoms makes such evaluation difficult. We therefore selected 85 asymptomatic individuals, from a cohort of individuals living in densely populated dormitories with active spread of SARS-CoV-2 infection (Fig 1A), who, based on their kinetics of appearance and disappearance of antibodies against NP and Spike, were likely exposed to SARS-CoV-2 at different timepoints. In these individuals, we measured directly ex vivo the quantity of IFN-γ producing T cells reactive to peptide pools covering different structural proteins (M, NP and Spike) and compared it to the T cell magnitude detected in symptomatic patients who were infected with SARS-CoV-2 at similar timepoints.

**Fig. 1.**
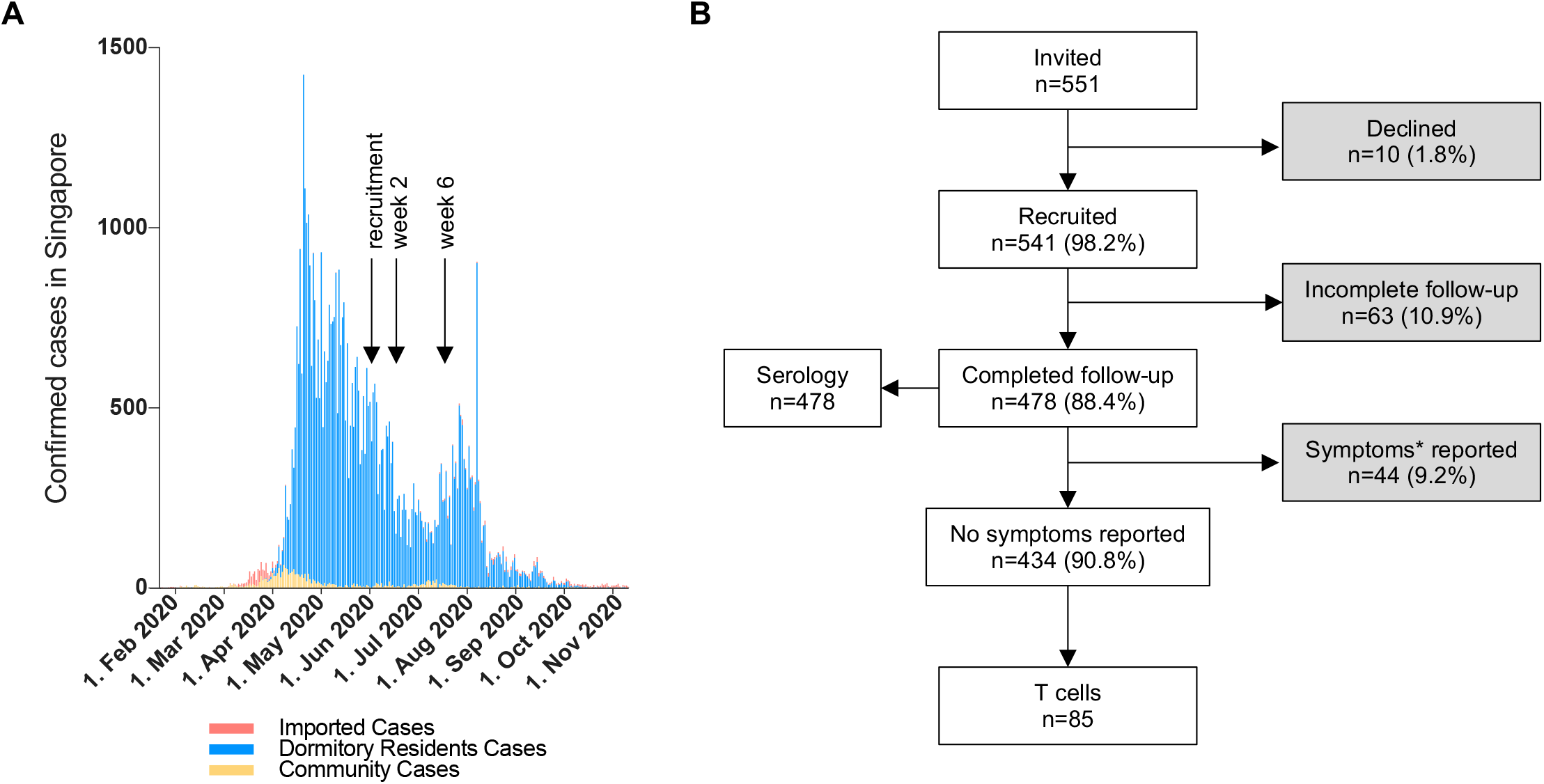
Enrolment of SARS-CoV-2 exposed donors. (**A**) Confirmed cases of SARS-CoV-2 infection in Singapore, divided into imported cases (red), cases among dormitory residents (blue) and all other community cases (yellow). The black arrows indicate the dates when blood samples were taken for the study. (**B**) Diagram showing the number of participants that were invited and recruited into the study. Participants who completed blood donations at recruitment and at 2- and 6-weeks follow-up were tested for serology of anti-NP IgG (Abbott) and sVNT-nAb (GenScript). An additional blood sample was taken at week 6 from 85 asymptomatic participants with distinct serological profiles for the analysis of SARS-CoV-2-specific T cell responses. *Symptoms: any of fever, cough, runny nose, sore throat, shortness of breath, fatigue, muscle ache, diarrhea or anosmia

We also evaluated, in both symptomatic and asymptomatic individuals, the functional profile of SARS-CoV-2-specific T cells. We designed a test to quantify secretion of IL-2, IFN-γ, IL-4, IL-6, IL-12p70, TNF-α, IL-1β and IL-10 directly in whole blood after T cell activation with peptide pools covering the different SARS-CoV-2 proteins. This experimental system not only measures the quantity of T cell cytokines (IL-2, IL-4, IFN-γ) directly secreted by SARS-CoV-2-specific T cells but can provide a direct evaluation of the T cells’ ability to activate inflammatory or regulatory pathways in other circulating immune cells. Our results provide experimental evidence that asymptomatic individuals mount a virus-specific T cell response that is indistinguishable from symptomatic patients in magnitude, but that is functionally more fit, being characterized by an augmented secretion of Th1 cytokines (IFN-γ and IL-2) associated with a proportionate and coordinated production of pro-(IL-6, TNF-α, IL-1β) and anti-inflammatory (IL-10) cytokines. The implications of these findings for pathology and vaccine designs are discussed.

## Results

### Longitudinal serology in asymptomatic SARS-CoV-2 infection

From April 2020, Singapore experienced large outbreaks of SARS-CoV-2 infections among migrant workers residing in densely populated dormitories (Fig 1A). In order to select asymptomatic individuals who were exposed to SARS-CoV-2 at different timepoints, we followed-up 478 residents of a SARS-CoV-2 affected dormitory, who donated blood at recruitment, and 2 and 6 weeks later for serological testing of anti-nucleoprotein (NP) IgG and neutralizing antibodies (nAb) (Fig 1B). At recruitment, 131 of the 478 (27.4%) participants were seropositive by either assay, with 6 (4.6%) reporting COVID-19 compatible symptoms in the preceding 4 weeks (Fig 2A). Over the 6-week follow-up, 171 of 347 (49.3%) initially seronegative individuals seroconverted by either assay, demonstrating ongoing exposure to SARS-CoV-2 infection. Only 15 (5.5%) individuals reported symptoms, generally mild, during the follow-up period. The majority of seropositive individuals (281/302; 93%) were asymptomatic. Individuals with symptoms were subsequently excluded from this study (Fig 1B).

**Fig. 2.**
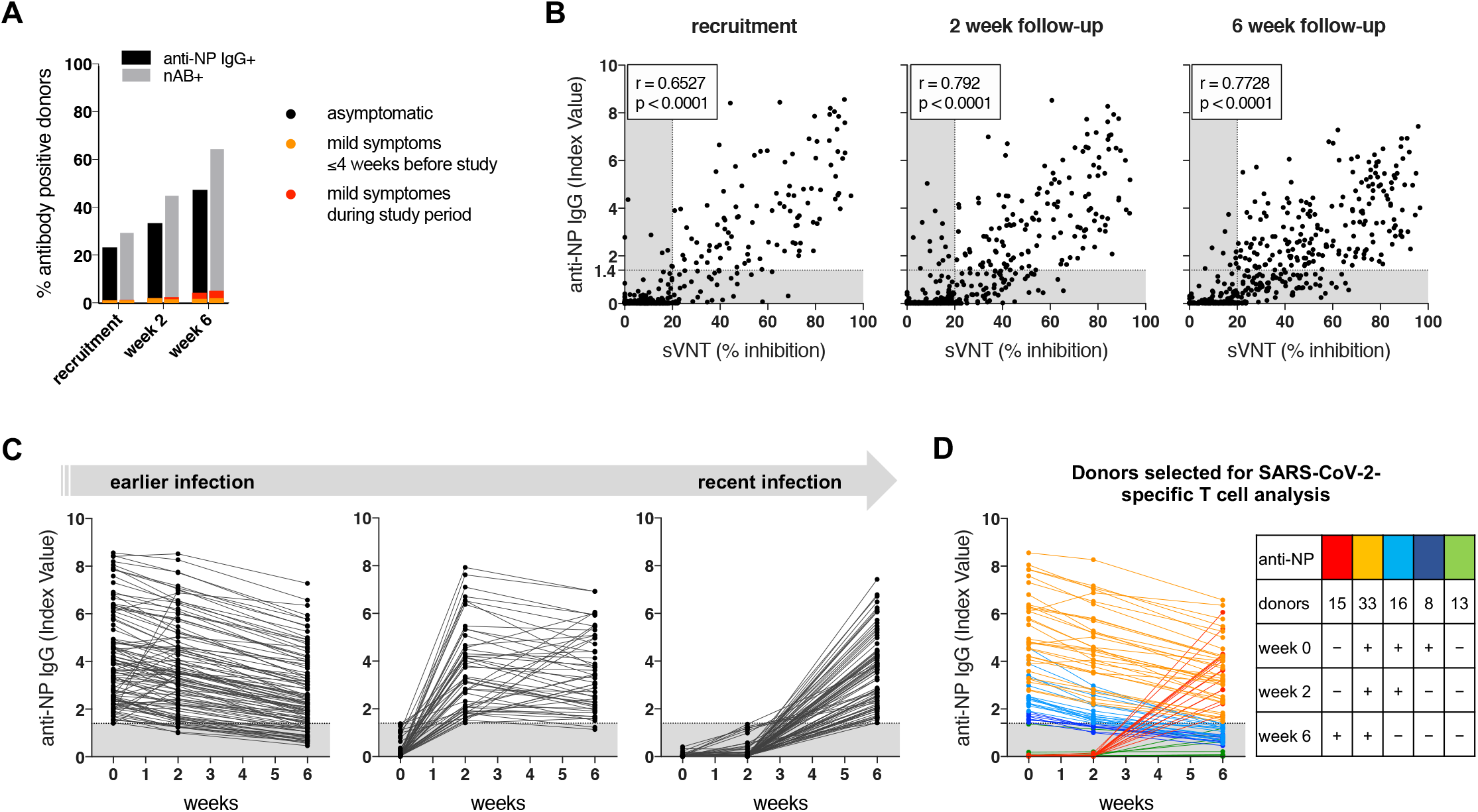
SARS-CoV-2-specific antibody profile shows increasing infection prevalence among the dormitory residents during the study period. (**A**) Percentage of donors positive for anti-NP IgG (black) and neutralizing antibodies (grey) at recruitment and after 2 and 6 weeks; antibody positive donors who experienced COVID-19 symptoms before (yellow) and during (red) the study are highlighted. (**B**) Dot plots show longitudinally the quantity of anti-NP IgG antibodies (y-axis) and the % inhibition by virus neutralization antibodies (sVNT; x-axis) in the serum of 434 asymptomatic study participants at recruitment (left), after 2 weeks (middle) and after 6 weeks (right). The grey area marks the limit of assay detection. Spearman correlation. (**C**) Longitudinal anti-NP IgG levels of asymptomatic donors who were seropositive at recruitment (left; n=106, left), who seroconverted at week 2 (n=52, middle) and who seroconverted by week 6 (n=77, right). (**D**) Anti-NP IgG serological profile of donors selected for SARS-CoV-2-specific T cell analysis at the 6-week timepoint. Donors with distinct antibody profiles are shown in different colors and are summarized in the table.

Kinetics of anti-NP IgG and of surrogate virus RBD neutralizing antibodies (sVNT-nAb) *(18)* in the asymptomatic study participants are shown in figure 2. Overall, there was a strong correlation between titers of anti-NP IgG and % of inhibition by sVNT-nAb (Fig 2B). However, at all timepoints, a higher percentage of asymptomatic donors was positive for sVNT-nAb (30.9%, 47.0% and 65.4% at recruitment, weeks 2 and 6, respectively) compared with anti-NP IgG (24.4%, 34.6% and 47.5%) (Fig 2A). Among the 106 individuals with positive anti-NP IgG levels at recruitment, 27 (25.5%) lost them during the six weeks (Fig 2B, C). In contrast, of the 134 participants who were seropositive for sVNT-nAb at baseline, only 12 (9.0%) became negative over the same time period (Fig S1).

### Virus-specific T cell quantification in asymptomatic SARS-CoV-2-exposed donors

To investigate SARS-CoV-2-specific T cells at the 6-week timepoint, which corresponds to 3 months after the first confirmed case in this dormitory, we selected, among the 434 asymptomatic study participants, 85 individuals with distinct antibody profiles (Fig 2C, D). Fifty-seven were anti-NP IgG positive at recruitment. Of these, 33 individuals remained persistently anti-NP IgG positive during the 6-week follow-up, while in 24 the antibody decreased below the limit of assay detection. We then selected 28 individuals who were seronegative at recruitment. Of these, 15 became anti-NP IgG positive, while 13 were persistently negative.

We first analyzed the frequency of cells reactive to three structural proteins of SARS-CoV-2 by ex vivo IFN-γ-ELISpot assays (Fig 3A). We designed pools of 15-mer peptides covering the whole NP and M proteins and a selected peptide pool of 15-mers covering the most T cell immunogenic regions of S (Fig 3A, Fig S2). As previously shown *(19)*, we demonstrate by intracellular cytokine staining that IFN-γ was produced by CD4 and CD8 T cells reactive to peptide stimulation, both directly ex-vivo and after expansion of short-term T cell lines re-stimulated with individual peptides (Fig S3).

**Fig. 3.**
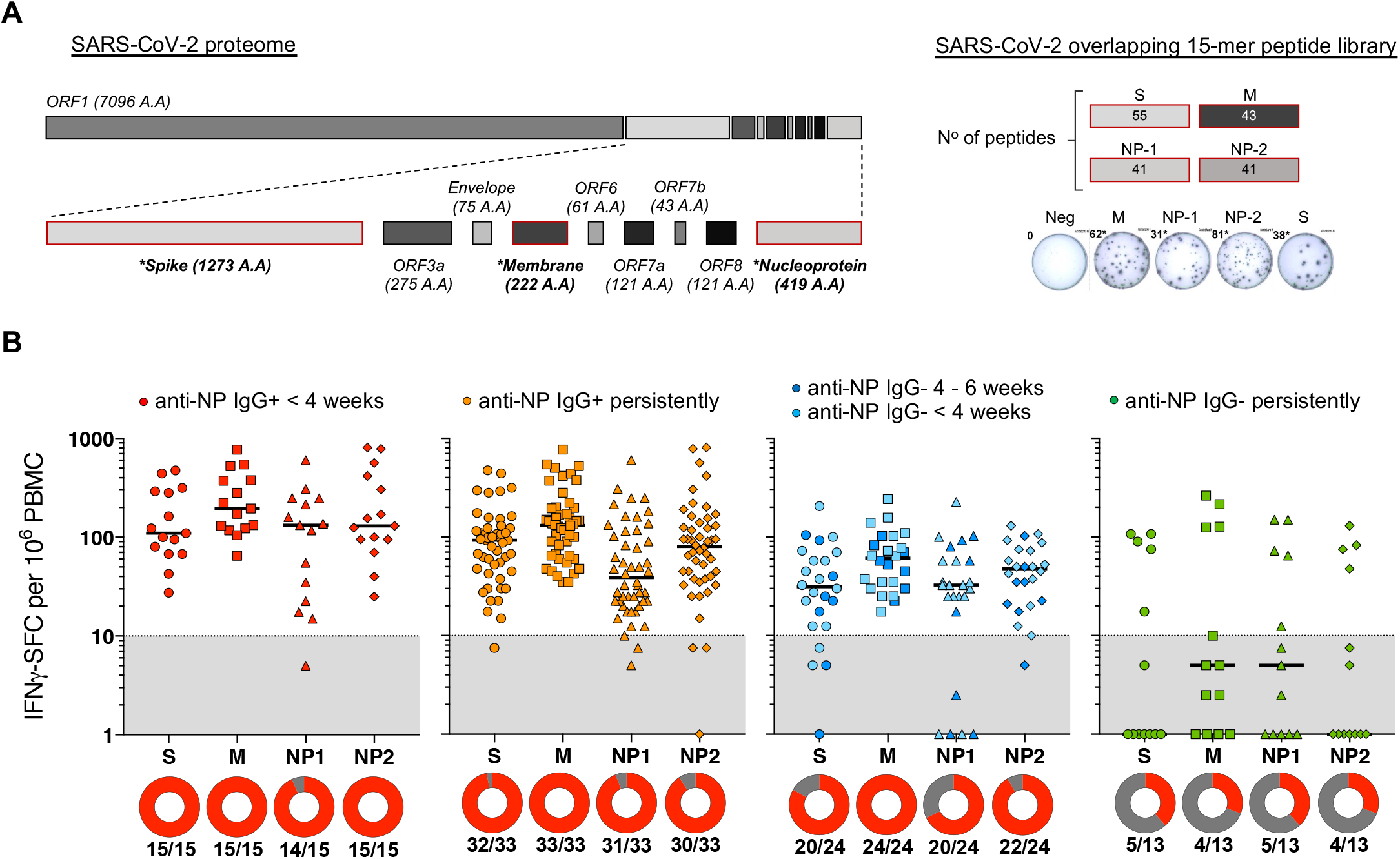
Frequency of T cells specific for different SARS-CoV-2 proteins in asymptomatic donors with distinct serological profiles. (**A**) SARS-CoV-2 proteome organization; analyzed proteins have a red outline and are marked by *. 15-mer peptides overlapping by 10 amino acids were split into pools covering nucleoprotein (NP1, NP-2) and membrane (M) and selected 15-mers covering the more T cell immunogenic regions of Spike (S). T cell-reactivity was tested by ex vivo IFN-γ-ELISpot. (**B**) The frequency of IFN-γ-spot forming cells (SFC) reactive to the individual peptide pools is shown for the asymptomatic donors with distinct serological profiles (line = median). IFN-γ-SFC ≥10/106 PBMC were considered positive (grey area is below limit of detection). Circles below represent the percentage of a positive response (red) to the individual peptide pools.

### SARS-CoV-2-specific T cells are present in all asymptomatic seropositive individuals

Cells reactive to SARS-CoV-2 peptide-pools were found in all anti-NP IgG positive individuals regardless of the duration of the antibody persistence (Fig 3B). Moreover, almost all anti-NP IgG positive individuals had SARS-CoV-2-specific T cells reactive to at least 3 pools simultaneously, except for 2 (of 24) individuals who had lost anti-NP IgG (Fig 3B). In all groups with different serology profiles, the M-peptide pool triggered a higher frequency of specific T cells in comparison to the S and the two NP pools.

Among the 13 persistently anti-NP IgG negative participants, 4 had SARS-CoV-2 specific T cells at a similar frequency as antibody positive participants (Fig 3B, right).

### Frequency, multi-specificity and kinetics of SARS-CoV-2-specific cellular immunity in symptomatic and asymptomatic infection

Next, we compared the SARS-CoV-2-specific T cell response of the asymptomatic cohort with the response in a cohort of hospitalized COVID-19 patients with mild to severe symptoms (Table S1). Within the first 3 months post viral exposure, the frequency of virus-specific T cells reactive to M, NP and S peptides was similar between symptomatic COVID-19 patients and asymptomatically-infected individuals, and both groups showed a higher frequency of M-pool-reactive T cells over other pools (Fig 4A). Importantly, nearly all COVID-19 patients and asymptomatic individuals with serological evidence of infection had T cells recognizing at least 3 peptide pools (Fig 4B).

**Fig. 4.**
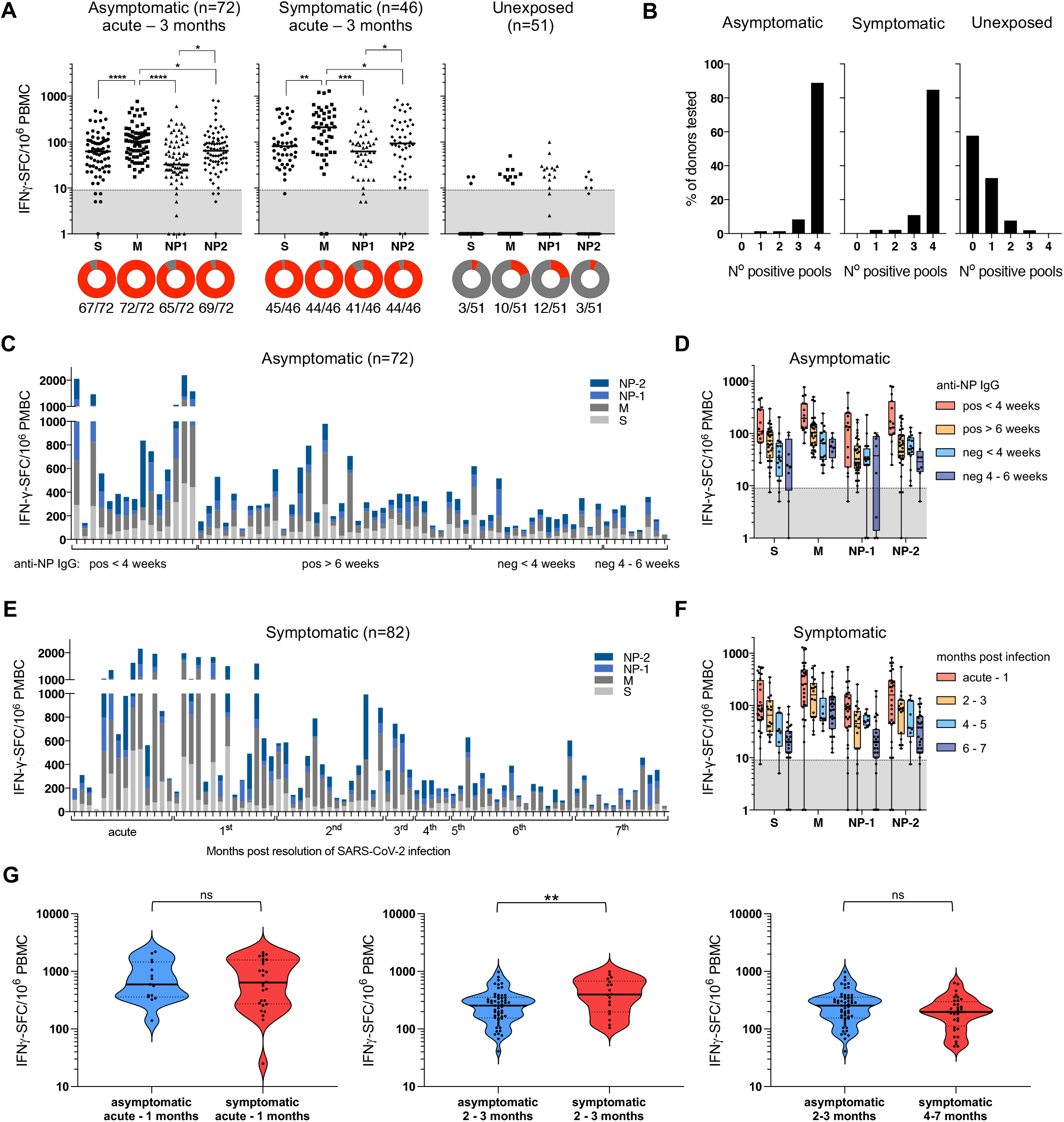
Dynamics and hierarchy of SARS-CoV-2-specific T cells in asymptomatic and symptomatic infection. (**A**) The frequency of IFN-γ-spot forming cells (SFC) reactive to the individual peptide pools is shown for asymptomatic SARS-CoV-2-infected donors who were serologically positive for anti-NP-IgG on at least one timepoint during the study period (n=72; left), for COVID-19 patients from acute to 3 months post infection (n=45; middle) and archived unexposed controls (n=51; right) (Line = median). Circles below represent the frequency of a positive (IFN-γ-SFC ≥10/10^6^ PBMC) response (red) to the individual peptide pools. RM one-way ANOVA with Greenhouse-Geisser correction, followed by Holm-Sidak’s multiple comparisons test. (**B**) Bar graphs show the percentage of donors reacting to the number of peptide pools tested. (**C**) Frequency of IFN-γ-SFC reactive to the individual peptide pools is shown for each donor of the asymptomatic cohort (n=72). Donors are organized according to their serological status of anti-NP IgG antibodies. (**D**) The frequency of IFN-γ-SFC reactive to the individual peptide pools is shown in asymptomatic donors grouped by anti-NP IgG status. (**E**) Frequency of IFN-γ-SFC reactive to the individual peptide pools is shown for symptomatic COVID-19 patients organized by months post clearance of SARS-CoV-2 infection (n=82). (**F**) The frequency of IFN-γ-SFC reactive to the individual peptide pools is shown in COVID-19 patients grouped by months post infection. (**G**) Frequency of SARS-CoV-2-peptide reactive cells in individuals with asymptomatic (blue) and symptomatic (red) infection during the acute phase till 1-month post recovery (left), in convalescents 2-3 months post infection (middle) and in asymptomatic at 2-3 months and in symptomatic at 4-7 months post infection (right). Mann-Whitney test.

In contrast, the pattern of cross-reactive SARS-CoV-2-specific T cells in archived samples from SARS-CoV-2 unexposed individuals was different. The overall frequency was lower (10 to 100 spots per 10^6^ PBMC) and where SARS-CoV-2-peptide reactive T cells were present (20 of 51) they mostly reacted to a single peptide pool (Fig 4A/B). Only 1 of 51 unexposed individuals tested had T cells specific for 3 peptide pools.

To investigate whether virus-specific T cells undergo clonal reduction over time, we compared their strength and peptide pool specificity in individuals who seroconverted to anti-NP IgG positive within the previous 4 weeks, individuals who seroconverted more than 6 weeks previously, and in initially seropositive individuals who became anti-NP IgG negative. The frequency of SARS-CoV-2-specific T cells was higher in recent seroconverters (anti-NP-IgG+ <4 weeks) than in those infected at an earlier time point (Fig 4C, D). Among those who became anti-NP IgG negative, the frequency of SARS-CoV-2-specific T cells was lower.

A decline in the frequency of SARS-CoV-2-specific T cells was also apparent from serial cross-sectional samples of COVID-19 patients from the acute phase over a period of 7 months post viral clearance (Fig 4E, F). Over time, some patients lost T cells specific to individual peptide pools, but M-specific T cells were maintained in all symptomatic and asymptomatic donors tested (Fig 4D, F).

Since there was a progressive reduction of SARS-CoV-2-specific T cells over time in all exposed individuals, we compared the kinetics of their decline between the asymptomatic and symptomatic cohorts. Among the recent seroconverters (<4 weeks), the frequency of SARS-CoV-2-specific T cells was comparable with that in symptomatic COVID-19 patients tested during infection and within 1 month post viral clearance (Fig 4G, left). However, the asymptomatic donors that had been exposed to SARS-CoV-2 about 2-3 months earlier (anti-NP+ >6 weeks earlier, first confirmed case 3 months earlier) showed a reduced frequency of SARS-CoV-2-specific T cells compared to patients who recovered from symptomatic infection 2-3 months earlier (Fig 4G, middle). Thus, the frequency of SARS-CoV-2-specific T cells declines faster after asymptomatic infection, and within 2-3 months reaches the levels detected in symptomatic patients found at 4-7 months post infection (Fig 4G, right).

### The pattern of cytokines induced by SARS-CoV-2-specific T cell activation in symptomatic and asymptomatic infection

In order to functionally define the antiviral cellular immune response present in asymptomatic and symptomatic SARS-CoV-2 infected individuals, we designed an assay where the level of Th1 (IFN-γ, IL-2, TNF-α), Th2 (IL-4), pro-inflammatory (IL-6, IL-1β, IL-12p70) and regulatory (IL-10) cytokines was tested directly in whole blood after overnight stimulation with SARS-CoV-2 peptide pools (Fig 5A). Cytokines secreted during overnight stimulation can either be produced by the activation of virus-specific T cells directly, or indirectly by other cell types that respond to the activation of antigen-specific T cells (Fig S4).

**Fig. 5.**
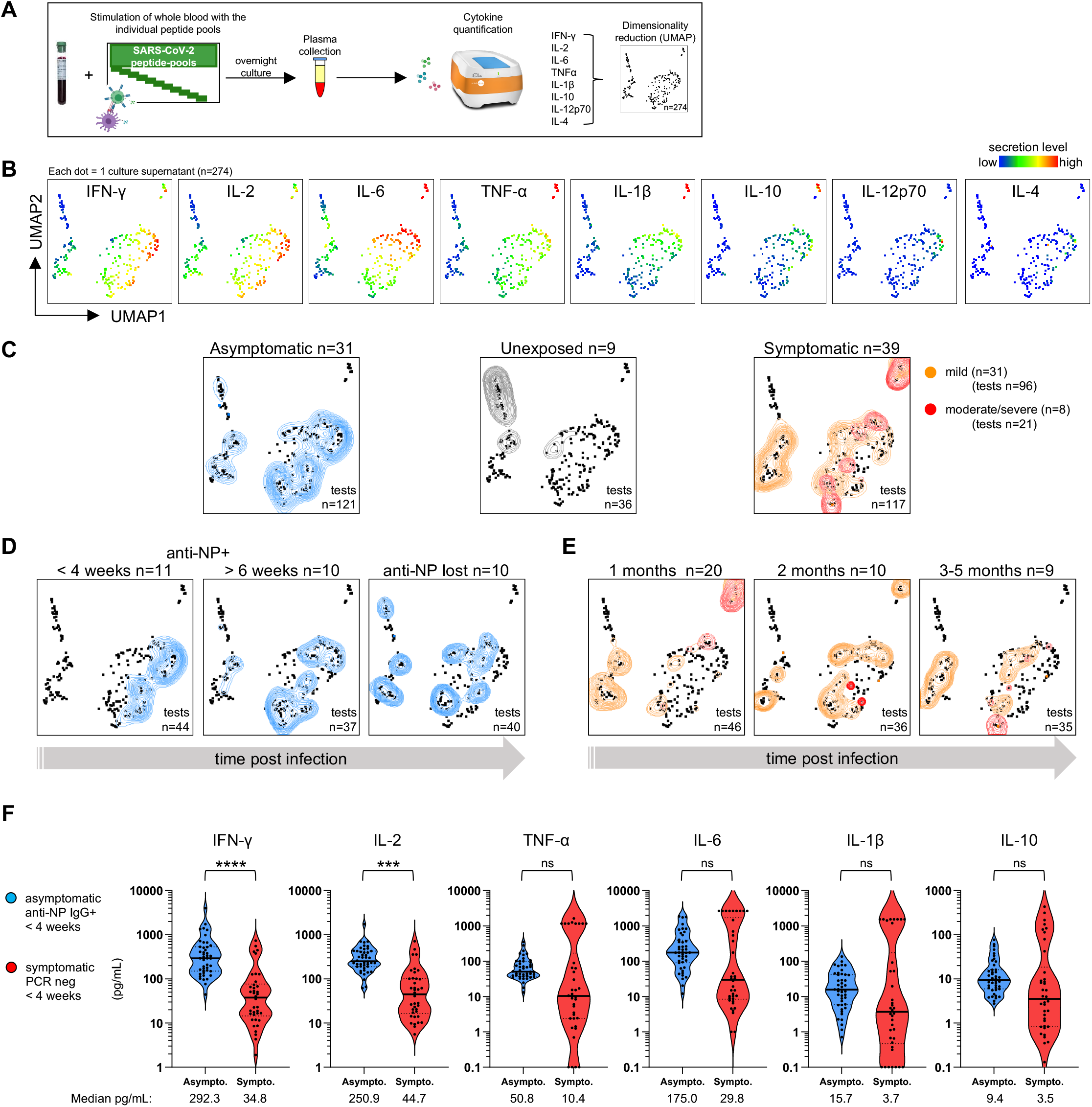
Cytokine secretion profile of whole blood from asymptomatic and symptomatic convalescents stimulated with SARS-CoV-2-peptide pools. (**A**) Schematic of whole blood stimulation with SARS-CoV-2-peptide pools overnight and analysis of the cytokine secretion profile (after DMSO control subtraction) using unsupervised clustering algorithm UMAP. (**B**) UMAP plots with cytokine secretion heatmaps. (**C**) Concatenated cytokine secretion profiles of peptide-pool stimulated whole blood from asymptomatic SARS-CoV-2 infected (left, blue, 31 donors, 121 tests), SARS-CoV-2 unexposed (middle, grey, 9 donors, 36 tests) and SARS-CoV-2 infected with mild (right, orange, 31 donors, 96 tests) and moderate to severe symptoms (right, red, 8 donors, 21 tests) overlaid on the global UMAP plot of all analyzed samples (black dots; each dot corresponds to one culture supernatant). (**D**) The cytokine secretion profile of the samples from the asymptomatic cohort separated by donors who seroconverted to anti-NP IgG+ <4 weeks ago (left), who were persistently seropositive (middle) and those who lost anti-NP IgG during the 6-week study period (right). (**E**) The cytokine secretion profile of the samples from the symptomatic cohort separated by donors who are in the 1st months (left), 2^nd^ months (middle) and 3^rd^ - 5^th^ months (right) post SARS-CoV-2 infection. (**F**) The amount of indicated cytokines secreted upon whole blood stimulation with the peptide pools is compared between donors who had recently (<4 weeks ago) an asymptomatic (blue) or a symptomatic (red) SARS-CoV-2 infection. Line = median concentration. Kruskal Wallis (non-parametric one-way ANOVA) followed by Dunn’s multiple comparisons test.

We used UMAP (Uniform Manifold Approximation and Projection) for unsupervised dimension reduction and clustering *(20)* of the secretoms of all peptide-stimulated samples after subtraction of cytokine levels present in corresponding DMSO controls (Fig 5B). This showed that secretion of cytokines classically produced by T cells, IFN-γ and IL-2, was overlapping. Moreover, IL-6 and IL-1β were co-secreted, as well as TNF-α and IL-10. IL-12p70 and IL-4 were undetectable in most samples. While stimulation of blood from unexposed donors did not induce any secretion of cytokines, the secretoms of SARS-CoV-2 infected donors differed between symptomatic and asymptomatic infection (Fig 5C). The majority of samples from asymptomatic donors co-secreted high levels of IFN-γ and IL-2, while the samples of symptomatic donors produced low levels of IFN-γ and IL-2. A subset of samples from symptomatic donors, enriched for those with severe disease, clustered with high levels of IL-6, TNF-α, IL-1β and IL-10 secretion (Fig 5C).

For the individuals with asymptomatic SARS-CoV-2 infection, we used the appearance, persistence and decline of anti-NP IgG levels as a proxy for time since infection. This distinguishes patients with recent infection (<4 weeks anti-NP IgG+) from those who were infected at earlier timepoints and allowed us to compare them with symptomatic patients that cleared the virus within the last 4 weeks. Strikingly, the cytokine profile between these two recently infected symptomatic and asymptomatic groups completely separated by UMAP (Fig 5D left, 5E left). Samples from recently infected asymptomatic individuals all co-secreted high levels of IFN-γ (median = 292 pg/mL) and IL-2 (median = 250 pg/mL), and intermediate levels of IL-6, TNF-α, IL-1β and IL-10. In contrast, samples from recently infected symptomatic donors co-secreted low levels of IFN-γ (median = 35 pg/mL) and IL-2 (median = 45 pg/mL), and a subset co-secreted very high levels of IL-6, TNF-α, IL-1β and IL-10 (Fig 5D, E, F).

Next, we correlated the cytokine levels secreted by peptide stimulation of whole blood with the frequency of IFN-γ spot forming cells detected by ELISpot assay in response to the same peptide pools (Fig 6). The secretion levels of IFN-γ and IL-2 induced by peptide stimulated samples correlated (p<0.0001) with the number of IFN-γ spots in both asymptomatic and symptomatic individuals, but higher levels of both cytokines in relation to the number of spots were detected in asymptomatic individuals.

**Fig. 6.**
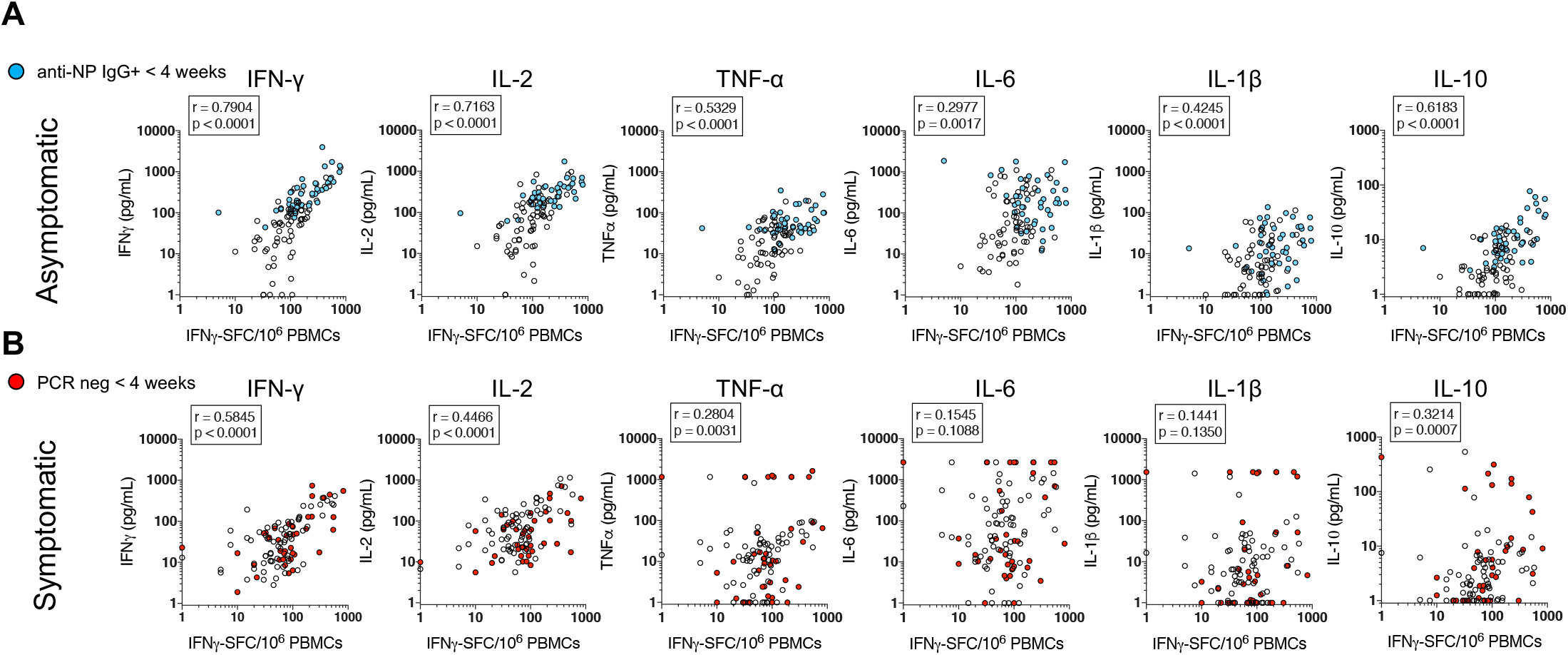
Frequency and function of SARS-CoV-2-peptide pool reactive cells. (**A**) The frequency of IFN-γ-spot-forming cells (SFC) reactive to the peptide pools analyzed by ELISpot assay (x-axis) is plotted against the amount of the cytokines secreted following whole blood stimulation with the identical peptide pools (y-axis) of the asymptomatic cohort. Samples from donors with the recent SARS-CoV-2 infection (<4 weeks anti-NP IgG+) are highlighted in blue. Spearman correlation. (**B**) The frequency of IFN-γ-secreting cells (SFC) reactive to the peptide pools analyzed by ELISpot assay (x-axis) is plotted against the amount of the cytokines secreted following whole blood stimulation with the identical peptide pools (y-axis) of the symptomatic cohort. Samples from donors with the recent SARS-CoV-2 infection (<4 weeks PCR neg) are highlighted in red. Spearman correlation.

In addition, levels of secreted pro-inflammatory (IL-6, TNF-α, IL-1β) and regulatory (IL-10) cytokines correlated strongly with the number of IFN-γ spots in asymptomatic individuals (Fig 6A). In some symptomatic patients, high levels of IL-6, TNF-α, IL-1β and IL-10 were produced but were uncorrelated with the number of spots detected in the ELISpot assays (Fig 6B).

## Discussion

Despite their ability to efficiently control the infection, asymptomatic individuals who clear SARS-CoV-2 have been hypothesized to mount a reduced antiviral adaptive immune response *(14)*. This hypothesis is supported mainly by measurement of SARS-CoV-2 specific antibodies *(14, 21, 22)* and B cell quantity *(23)*.

Our study shows clearly that the ability to mount a significant virus-specific T cell response is not necessarily associated with symptom severity. Our results demonstrate that the overall magnitude of T cell responses against different structural proteins was similar in both asymptomatic individuals and COVID-19 patients. Moreover, the T cells induced by an asymptomatic infection appear to secrete higher quantities of IFN-γ and IL-2 and trigger a more coordinated production of pro-inflammatory and regulatory cytokines than T cells of symptomatic COVID-19 patients. Overall, we conclude that asymptomatic SARS-CoV-2 infected individuals are able to raise an efficient and balanced anti-viral cellular immunity which protects the host without causing any apparent pathology.

One of the principal characteristics of our study was in the ability to select, through a 6-week longitudinal serological study, asymptomatic individuals in whom the time of viral exposure could be reasonably estimated. The first confirmed SARS-CoV-2 infection in this dormitory was identified in mid-April 2020, so asymptomatic, seropositive individuals in our cohort were unlikely to have been infected more than 3 months before sampling (performed in mid-July 2020) (Fig 1A). More importantly, by evaluating the period of SARS-CoV-2 antibody seroconversion, we could separate individuals who were likely infected more than 6 weeks before sampling from those who were likely infected within the last 4 weeks.

The ability to estimate the time of infection allowed us to compare the magnitude and function of SARS-CoV-2-specific T cells present in asymptomatically-infected individuals with those of COVID-19 patients at similar timepoints during and after infection. We demonstrated that all asymptomatic individuals capable of producing anti-SARS-CoV-2 antibodies also mount a T cell response that simultaneously targets different structural proteins of SARS-CoV-2 (NP, M and S). In addition, we observed that, similarly to COVID-19 recovered patients, SARS-CoV-2-specific T cell frequency in asymptomatic individuals declined progressively over time. Surprisingly, more recently infected asymptomatic individuals possess a SARS-CoV-2-specific T cell frequency of indistinguishable magnitude to the one detected in COVID-19 patients (mild, moderate and severe) tested during acute infection (SARS-CoV-2 PCR positive) or within 1 month after viral clearance. This finding further supports the accumulating evidence that the quantity of SARS-CoV-2-specific T cells is not proportional to disease severity during the early phase of infection *(9, 10)*. This is in contrast to the positive correlation between SARS-CoV-2-specific T cell frequency and presence/severity of symptoms reported by others. In those studies, however, SARS-CoV-2-specific T cell responses were evaluated several months after recovery *(15, 16, 24, 25)*.

We further show that despite a similar magnitude of virus-specific T cells induced in asymptomatic and symptomatic infection, SARS-CoV-2-specific T cells seem to decline faster in asymptomatic individuals. SARS-CoV-2-specific T cell frequency detected 2-3 months after asymptomatic infection was lower than that detected in COVID-19 patients at similar time after infection and identical to what was detected in COVID-19 recovered individuals 6 months after infection. A more extended study of the persistence of SARS-CoV-2-specific T cells in our asymptomatic cohort is needed to confirm such a trend. Nevertheless, at this moment our data suggest that the recently reported higher frequency of SARS-CoV-2-specific T cells in COVID-19 recovered patients in comparison to asymptomatic individuals 6 months after recovery, might not be caused by a higher set point present during the early phases of infection as suggested *(24)*. An alternative explanation could be that SARS-CoV-2-specific T cells persist longer at a higher frequency in COVID-19 recovered patients, since viral antigen might persist more in symptomatic patients who usually have higher quantity of viral replication *(26)*. The recent observation that SARS-CoV-2-specific B cells progressively evolved after COVID-19 recovery due to the presence of SARS-CoV-2 antigen in the small bowel *(27)*, makes such hypothesis even more plausible.

Finally, we also show that asymptomatic SARS-CoV-2 infected individuals mount an efficient anti-viral cellular immunity, by analyzing the quantity of cytokines secreted in whole blood after pulsing with different SARS-CoV-2 peptide pools and by correlating this quantity with the number of spots detected in our ELISpot analysis. We utilized this relatively simple approach for several reasons. In our study, we wanted to prioritize a direct ex vivo analysis of the T cell response in fresh blood rather than in frozen PBMC, since freezing and thawing can reduce the detection of CD4 T cell responses *(28)* that represent the majority of virus-specific T cells in SARS-CoV-2 infection *(19, 29)*. We also wanted to analyze the global impact that T cell activation might have on other immune cells, particularly of the myeloid compartment, which is known to contribute substantially to the inflammatory reactions associated with SARS-CoV-2 infection *(3, 30, 31)*. Clearly, this whole blood cytokine assay has limitations, since it does not enable distinction of the cellular source of the different cytokines. However, it provides a holistic snapshot of the virus-specific cellular immunity and can be performed in a large group of asymptomatic individuals tested in a limited time. The results obtained with unsupervised clustering were extremely clear in describing distinct cytokine secretion patterns in symptomatic versus asymptomatic individuals and revealed possible features of SARS-CoV-2-specific T cell responses that are worthy of more detailed characterization in future studies.

We observed a high secretion of TNF-α, IL-6, IL-1β and IL-10 exclusively in the blood of a subset of patients who recently recovered from COVID-19, especially in patients with severe COVID-19. The high secretion of pro-inflammatory cytokines was, however, neither proportional to the quantity of IFN-γ and IL-2 detected in the same assay nor to the number of IFN-γ spots detected in the same patients analyzed in parallel with ELISpot. These findings seem to confirm that myeloid cells present in patients who recovered from symptomatic COVID-19 disease have been reprogrammed to promote inflammatory signals and are hyper-responsive to IFNs *(30)*.

Secretion of the pro-inflammatory cytokines IL-6, TNF-α, IL-1β was also detected in peptide-stimulated blood of asymptomatic individuals. Yet, their quantity was directly proportional to the quantity of T cell cytokines (IL-2, IFN-γ). Surprisingly, the quantity of IL-2 and IFN-γ was much higher in asymptomatic individuals than in COVID-19 recovered patients. Moreover, the close correlation of IFN-γ spots and cytokine production detected in the same experiments strongly suggests that T cells of asymptomatic individuals are endowed with a higher IFN-γ and IL-2 production capacity than the T cells of COVID-19 recovered patients.

To our knowledge, these are the first data to show, through direct comparison, that SARS-CoV-2-specific cellular immune responses are more efficient in asymptomatic than symptomatic individuals. Previous results are supporting our findings: a low IFN-γ production of total *(32)* and SARS-CoV-2-specific T cells *(33)* has been reported in COVID-19 patients, while conversely, high quantity of IFN-γ RNA was found in CD4 T cells of asymptomatic individuals *(17)*.

Finally, we detected production of IL-10 proportional to T cell cytokines (IL-2 and IFN-γ) and to the numbers of spots activated by similar SARS-CoV-2 peptide activation only in asymptomatic individuals. Unfortunately, technical challenges (low quantity of IL-10 secretion by T cells, lack of fresh blood of asymptomatic subjects recently infected) has so far precluded the direct visualization of IL-10 production by T cells and not by other cell types (B cells, monocytes) known to secrete IL-10 *(34)*. In conclusion, the overall picture of SARS-CoV-2-specific cellular immune responses detected by a combination of two different functional assays shows that patients with asymptomatic infection mount a coordinated and balanced activation of virus-specific T cells able to trigger a production of high quantities of IFN-γ and IL-2 linked with secretion of IL-10. Animal models have already shown that the ability of T cells to secrete IFN-γ and IL-10 simultaneously led to an effective viral control without triggering pathological processes *(12, 35)*. We suggest that this could be the functional signature of protective virus-specific cellular immune responses in asymptomatic SARS-CoV-2 infection. A detailed analysis at single cell level of the functional profile of SARS-CoV-2-specific T cells in symptomatically and asymptomatically infected individuals will be needed to formally demonstrate such hypothesis.

## Materials and Methods

### Ethics statement

The study of migrant workers was approved by the Singapore Ministry of Health under Section 59A of the Infectious Diseases Act (2015), thus institutional review board approval was not required and all participants gave verbal consent to participate. All COVID-19 patients provided written informed consent and their study was approved by the Institutional Review Boards of NUS (H-20-006), SingHealth (CIRB/F/2018/2387 and CIRB/F/2018/3045) and the National Healthcare Group Domain Specific Review Board (2012/00917).

### Study population

The asymptomatic cohort comprised 541 men aged 19-59 years (median 35 years) recruited from randomly-selected rooms within a SARS-CoV-2 affected dormitory housing over 4000 migrant workers and followed up prospectively from May to July 2020. Of these, 478 (88.4%) participants provided blood samples at recruitment, and at 2 and 6 weeks (Fig 1). Participants also declared symptoms at the time of blood draws and their information was cross-checked with the medical post based in the dormitory. At the 6-week follow-up, a subset of 85 asymptomatic participants provided an additional blood sample for assessment of cell-mediated immune responses. Participants were not tested for infection by polymerase chain reaction (PCR), because at the time of the study, testing of asymptomatic individuals without clinical indication was not recommended.

For hospitalized symptomatic patients, blood samples were obtained from the PCR positivity phase till 193 days post PCR negativity from recovered COVID-19 patients (n=76; n=16 were sampled longitudinally; Table S1). All archived healthy donors’ samples were collected before June 2019.

### Antibody quantification

Sera were tested for anti-nucleoprotein (NP) IgG antibodies (CMIA, Abbott Laboratories) on an Abbott Architect i2000SR automated instrument; a signal to cutoff ratio of ≥1.4 was defined as a positive result following the manufacturer’s recommendation. A second serum aliquot was tested with a surrogate virus neutralization test that detects isotype-independent RBD neutralizing antibodies (cPass, GenScript) *(18)*. Based on the manufacturer’s instructions, a virus inhibition threshold of ≥20% was considered a positive result.

### SARS-CoV-2-specific T cell quantification

SARS-CoV-2-specific T cells were tested as described before *(19)*. Briefly, peripheral blood mononuclear cells (PBMC) were isolated using Ficoll-Paque and directly tested by IFN-γ-ELISpot for reactivity to 4 SARS-CoV-2 peptide pools of 15-mers covering NP (NP-1, NP-2) and membrane (M), and one pool of 55 peptides covering the most immunogenic regions of Spike (S) (Fig S2). ELISpot plates (Millipore) were coated with human IFN-γ antibody (1-D1K, Mabtech) overnight at 4ºC. 400,000 PBMC were seeded per well and stimulated for 18h with pools of SARS-CoV-2 peptides (2 μg/ml). For stimulation with peptide matrix pools or single peptides, a concentration of 5 μg/ml was used. Subsequently, the plates were developed with human biotinylated IFN-γ detection antibody (7-B6-1, Mabtech), followed by incubation with Streptavidin-AP (Mabtech) and KPL BCIP/NBT Phosphatase Substrate (SeraCare). To quantify positive peptide-specific responses, 2x mean spots of the unstimulated wells were subtracted from the peptide-stimulated wells, and the results expressed as spot forming cells (SFC)/10^6^ PBMC. We excluded the results if negative control wells had >30 SFC/10^6^ PBMC or positive control wells (PMA/Ionomycin) were negative.

### Cell culture for T cell expansion

T cell lines were generated as follows: 20% of PBMCs were pulsed with 10 μg/ml of the overlapping SARS-CoV-2 peptides for 1 hour at 37°C, subsequently washed, and cocultured with the remaining cells in AIM-V medium (Gibco; Thermo Fisher Scientific) supplemented with 2% AB human serum (Gibco; Thermo Fisher Scientific). T cell lines were cultured for 10 days in the presence of 20 U/ml of recombinant IL-2 (R&D Systems).

### Flow cytometry

PBMC or expanded T cell lines were stimulated for 5h (or 7h) at 37°C with or without SARS-CoV-2 peptide pools (2 μg/ml). After 1h (or 3h,) 10 μg/ml brefeldin A (Sigma-Aldrich) and 1× monensin (Biolegend) were added. Cells were stained with the yellow LIVE/DEAD fixable dead cell stain kit (Invitrogen) and surface marker: anti-CD3 (SK7 or OKT3, Biolegend), anti-CD4 (SK3), anti-CD8 (SK1), anti-CD14 (TUK4, Miltenyi Biotec), anti-CD16 (3G8, Biolegend), anti-CD19 (SJ25-C1), anti-CD56 (HCD56, Biolegend). Cells were subsequently fixed and permeabilized using the Cytofix/Cytoperm kit (BD Biosciences-Pharmingen) and stained with anti-IFN-γ (25723, R&D Systems), anti-IL-2 (MQ1-17H12), anti-IL-6 (MQ2-13A5), anti-IL-10 (JES3-19F1) and anti-TNF-α (MAb11) antibodies and analyzed on a BD-LSR II FACS Scan. Data were analyzed by FlowJo (BD Biosciences). Antibodies were purchased from BD Biosciences-Pharmingen unless otherwise stated.

### Whole blood culture with SARS-CoV-2 peptide pools

320 μl of whole blood drawn on the same day were mixed with 80 μl RPMI and stimulated with pools of SARS-CoV-2 peptides (M, NP1, NP2 or S; 2 μg/ml) or a DMSO control. After 15 hours of culture, the culture supernatant (plasma) was collected and stored at −80ºC until quantification of cytokines.

### Cytokine quantification and analysis

Cytokine concentrations in the plasma were quantified using an Ella machine with microfluidic multiplex cartridges measuring IFN-γ, IL-2, TNF-α, IL-4, IL-6, IL-1β, IL-12p70 and IL-10 following the manufacturer’s instructions (ProteinSimple). The level of cytokines present in the plasma of DMSO controls was subtracted from the corresponding peptide pool stimulated samples. Subsequently, concentrations of each cytokine in all culture supernatants were transformed using the logicle transformation function and UMAP was run using 15 nearest neighbors (*nn*), a *min_dist* of 0.2 and euclidean distance *(20)*. The results obtained from UMAP analyses were incorporated as additional parameters and converted to .fcs files, which were then loaded into FlowJo to generate heatmaps of cytokine secretion on the reduced dimensions.

### Statistical analysis

All tests are stated in the figure legends. P-values (all two-tailed): <0.5 = *; <0.01 = **; <0.001 = ***; <0.0001 = ****.

## Acknowledgements

We thank Xin Mei Ong, Wan Rong Sia, Madeline Kwek and Charles Tiu for performing the surrogate virus neutralization testing in this study.

## Author contributions

NLB, ATT and AB designed the experiments; HEC and LYH conceived and contributed to study design for the migrant workers study; WNC, BLL, LFW, SHL, WYW performed antibody analysis; CYLT, KK, JMEL and AC performed all other experiments and analyzed data; JML, LT, NS, MZ, ZMT, VK, LJS contributed to design and implementation of the migrant workers study; NLB, CAD and AB analyzed and interpreted the data and prepared the figures; NLB and AB wrote the paper; YJT, PAT, SK, DL, JGHL recruited COVID-19 patients and provided all clinical data. AB designed and coordinated the study; CCT conceived and led the migrant workers study.

## Supplementary Materials

**Fig. S1.**
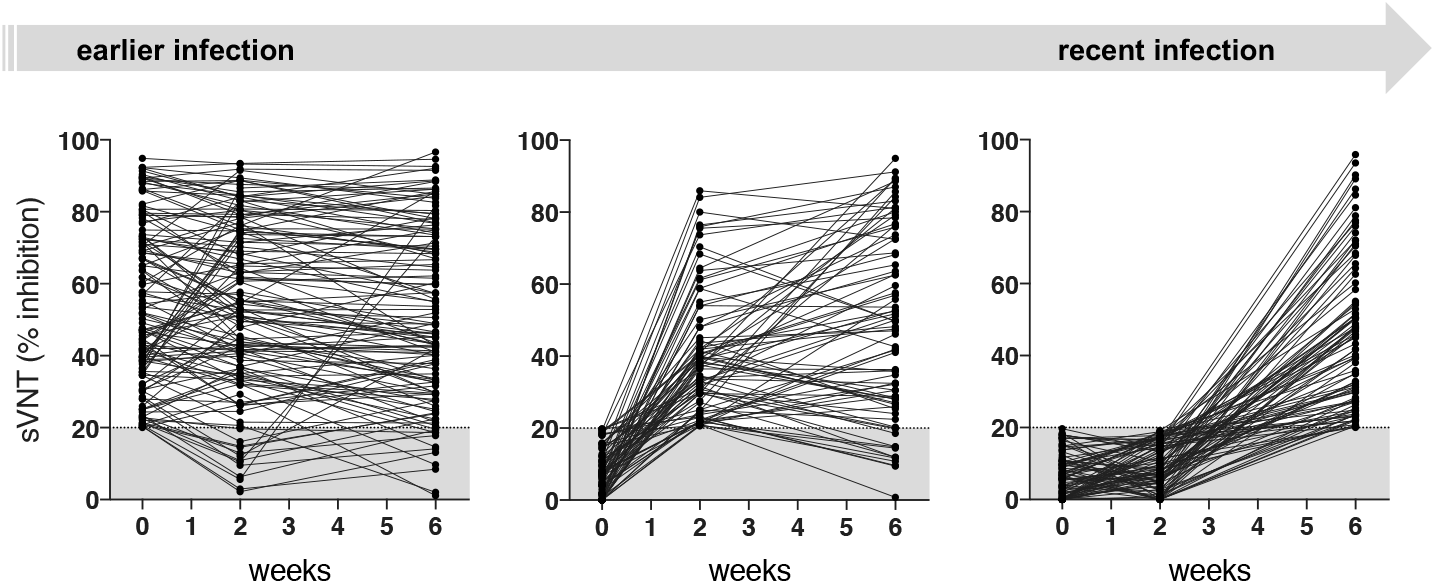
Kinetics of SARS-CoV-2 neutralizing antibody profile in the asymptomatic study cohort. Longitudinal levels of neutralizing antibodies measured as % inhibition by surrogate virus neutralization test (sVNT) in asymptomatic donors who were seropositive at recruitment (left; n=134, left), who seroconverted at week 2 (n=81, middle) and who seroconverted by week 6 (n=93, right). The grey area marks the limit of assay detection.

**Fig. S2.**
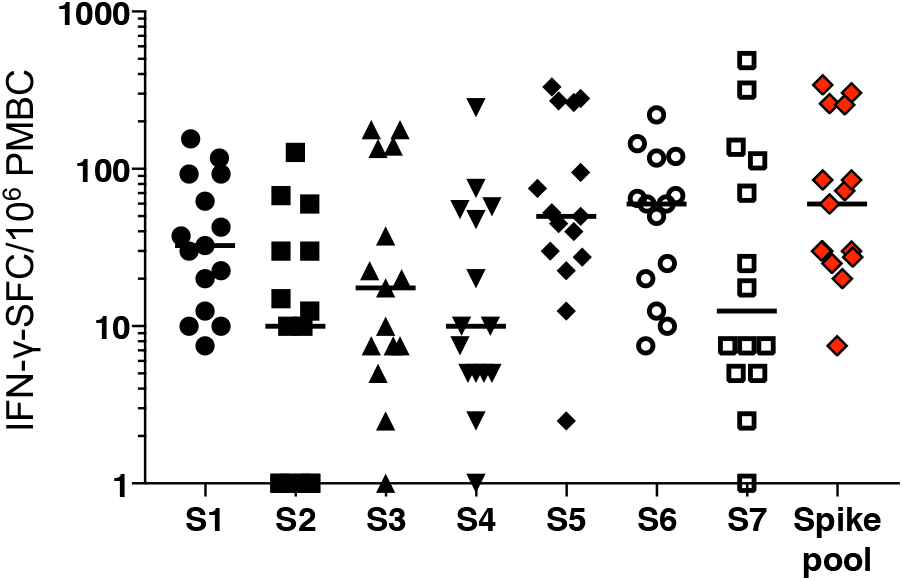
T cells reactive to different pools of Spike peptides in COVID-19 convalescents. Spike is a long protein with 1276 amino acids, thus it requires 253 15-mer peptides overlapping by 10 amino acids to cover the whole protein, thus 7 pools of about 40 peptides. To reduce the number of peptides pools to test, we selected a single “Spike pool” comprised of 55 peptides. For the selection, all sequences of published SARS-CoV-1 epitopes (www.iedb.org; positive assays only, T cells assays, host: human) were aligned with the library of Spike-SARS-CoV-2 15-mers. We selected the 15-mer peptides that cover the homologue sequence of the described SARS-CoV-1 epitope sequences. In addition, we added the 15-mer peptides that cover the predicted SARS-CoV-2 Spike epitopes published by Grifoni et al *(36)*. The 55 peptides cover 40.5% of the Spike protein. The frequency of reactive cells to the selected Spike pool (red) was compared to the 7 pools of 15-mers overlapping by 10 amino acids covering together the entire Spike protein (S1-S7) in 15 COVID-19 convalescents.

**Fig. S3.**
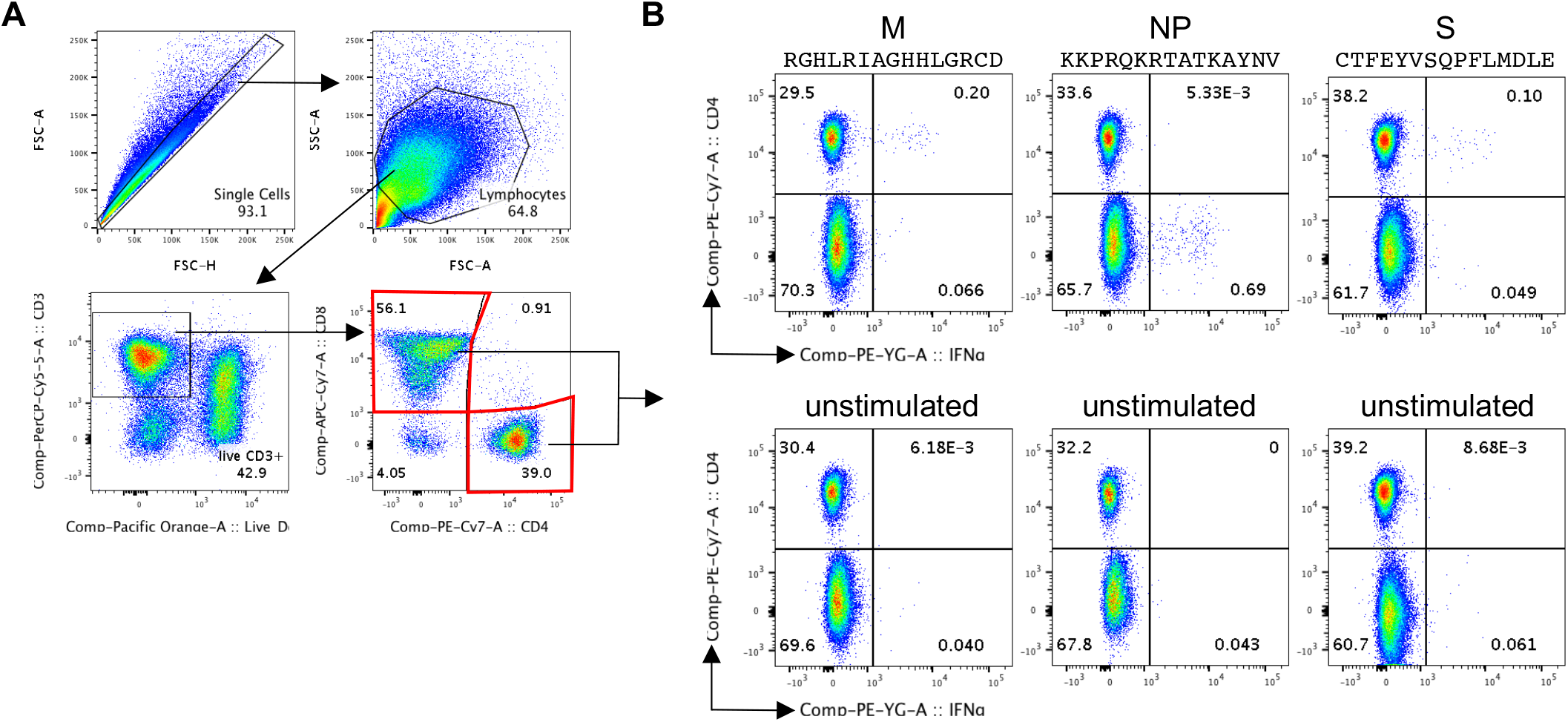
CD4+ and CD8+ T cells are reactive to stimulation with SARS-CoV-2-specific peptides. For 3 asymptomatic donors (<4 weeks anti-NP IgG+), short-term T cell lines were generated by PBMC stimulation with the different peptide pools and a 10-day expansion protocol *(19)*. A peptide pool-matrix strategy was applied which identified single T cell epitopes. Subsequently, the short-term T cell lines were re-stimulated for 5h with the identified single peptides and analyzed by intracellular cytokine staining for IFN-γ. (**A**) The gating strategy to identify CD4+ and CD8+ T cells is shown. (**B**) Dot plots show examples of CD4+ and CD8+ T cells producing IFN-γ in response to stimulation with three different peptides (upper panels) and the corresponding unstimulated controls (lower panels).

**Fig. S4.**
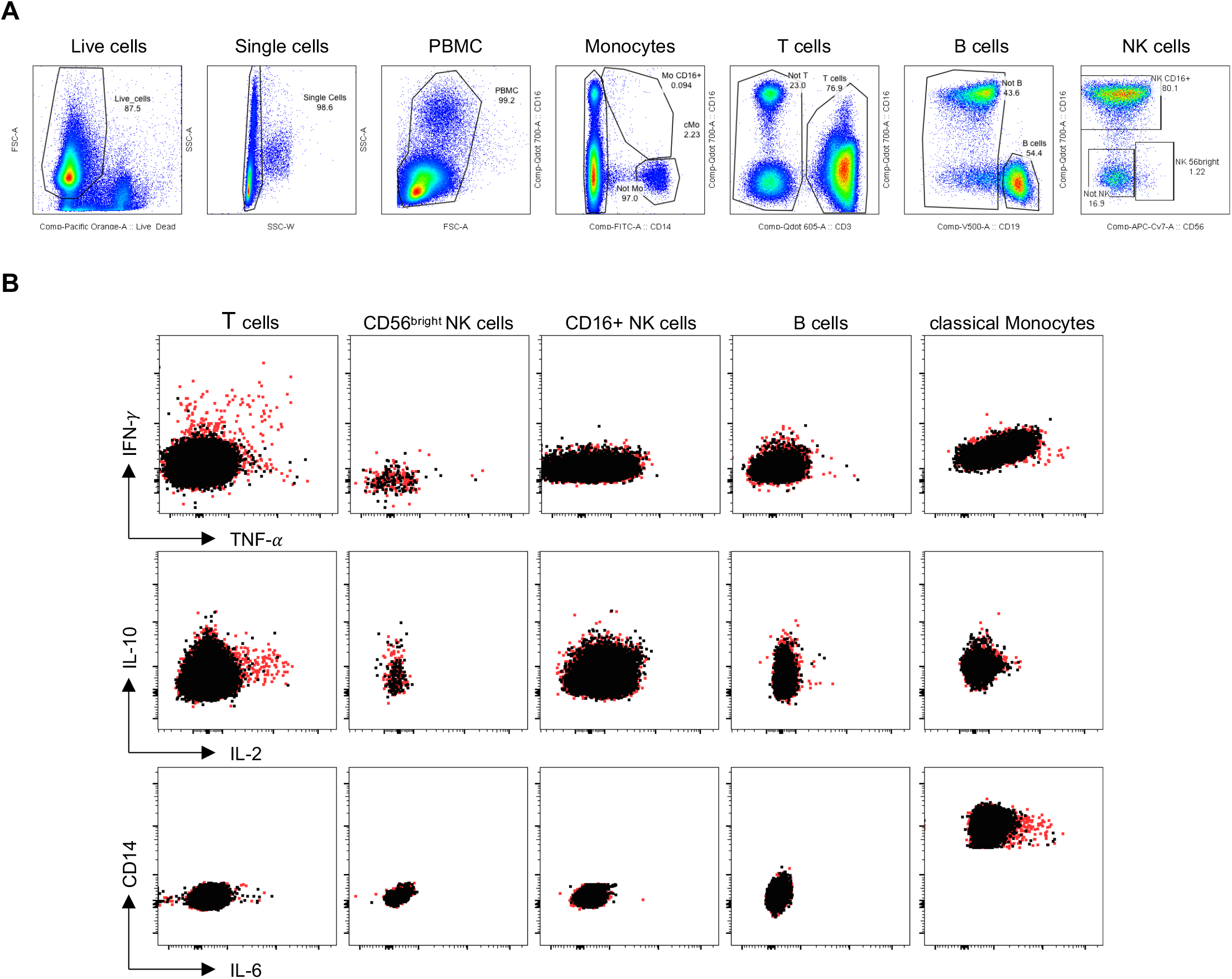
Cytokines produced by immune cells following stimulation with SARS-CoV-2 peptide pools. PBMC were stimulated for 5h (or 7h for IL-6) with SARS-CoV-2 peptide pools and analyzed by intracellular cytokine staining. (**A**) The gating strategy to identify the different immune cell subsets is shown. (**B**) Dot plots show examples of T cells producing IFN-γ, TNF-a, IL-2 and monocytes producing IL-6 in response to stimulation with the M peptide pool (red) overlaid with the corresponding unstimulated controls (black).

**Table S1:**
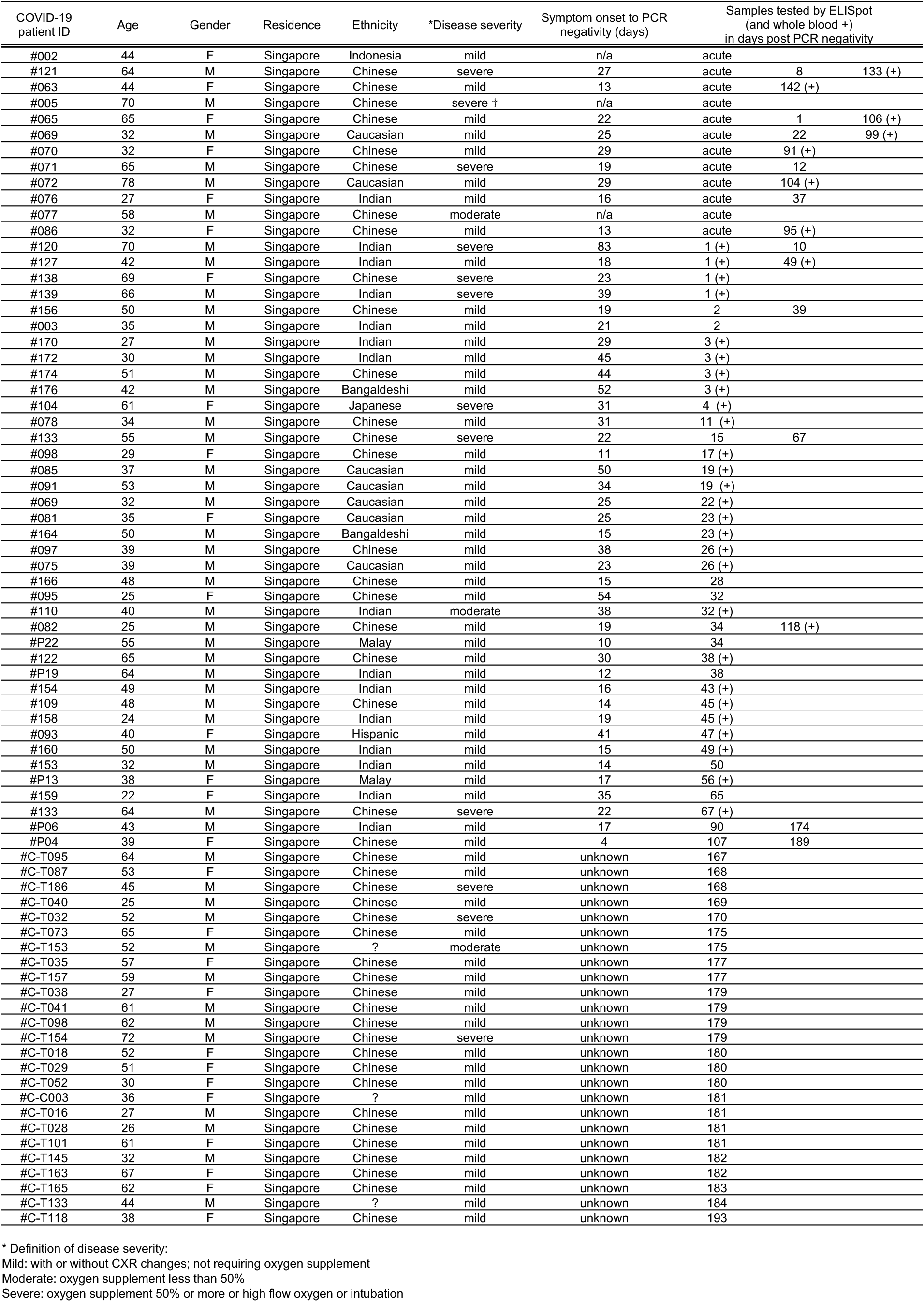
Cohort of symptomatic COVID-19 patients. SARS-CoV-2-specific T cells for S, M, NP-1 and NP-2 were analyzed in 76 COVID-19 patients by ELISpot assay. From 16 patients, T cells were tested longitudinally at 2-3 different timepoints post PCR negativity. In addition, whole blood was stimulated with the peptide pools from 38 donors, 1 donor was tested on two different time points. The tested samples are labelled with +.

## References

1. S. A. Vardhana, J. D. Wolchok, The many faces of the anti-COVID immune response, J. Exp. Med. 217, 166–10 (2020).

2. L. Kuri-Cervantes, M. B. Pampena, W. Meng, A. M. Rosenfeld, C. A. G. Ittner, A. R. Weisman, R. S. Agyekum, D. Mathew, A. E. Baxter, L. A. Vella, O. Kuthuru, S. A. Apostolidis, L. Bershaw, J. Dougherty, A. R. Greenplate, A. Pattekar, J. Kim, N. Han, S. Gouma, M. E. Weirick, C. P. Arevalo, M. J. Bolton, E. C. Goodwin, E. M. Anderson, S. E. Hensley, T. K. Jones, N. S. Mangalmurti, E. T. Luning Prak, E. J. Wherry, N. J. Meyer, M. R. Betts, Comprehensive mapping of immune perturbations associated with severe COVID-19, Science Immunology 5, eabd7114–20 (2020).

3. A. Silvin, N. Chapuis, G. Dunsmore, A.-G. Goubet, A. Dubuisson, L. Derosa, C. Almire, C. Hénon, O. Kosmider, N. Droin, P. Rameau, C. Catelain, A. Alfaro, C. Dussiau, C. Friedrich, E. Sourdeau, N. Marin, T.-A. Szwebel, D. Cantin, L. Mouthon, D. Borderie, M. Deloger, D. Bredel, S. Mouraud, D. Drubay, M. Andrieu, A.-S. Lhonneur, V. Saada, A. Stoclin, C. Willekens, F. Pommeret, F. Griscelli, L. G. Ng, Z. Zhang, P. Bost, I. Amit, F. Barlesi, A. Marabelle, F. Pène, B. Gachot, F. André, L. Zitvogel, F. Ginhoux, M. Fontenay, E. Solary, Elevated Calprotectin and Abnormal Myeloid Cell Subsets Discriminate Severe from Mild COVID-19, Cell 182, 1401–1418.e18 (2020).

4. R. Nienhold, Y. Ciani, V. H. Koelzer, A. Tzankov, J. D. Haslbauer, T. Menter, N. Schwab, M. Henkel, A. Frank, V. Zsikla, N. Willi, W. Kempf, T. Hoyler, M. Barbareschi, H. Moch, M. Tolnay, G. Cathomas, F. Demichelis, T. Junt, K. D. Mertz, Two distinct immunopathological profiles in autopsy lungs of COVID-19, Nature Communications 11, 5086–13 (2020).

5. A. G. Laing, A. Lorenc, I. del Molino del Barrio, A. Das, M. Fish, L. Monin, M. Muñoz-Ruiz, D. R. McKenzie, T. S. Hayday, I. Francos-Quijorna, S. Kamdar, M. Joseph, D. Davies, R. Davis, A. Jennings, I. Zlatareva, P. Vantourout, Y. Wu, V. Sofra, F. Cano, M. Greco, E. Theodoridis, J. Freedman, S. Gee, J. N. E. Chan, S. Ryan, E. Bugallo-Blanco, P. Peterson, K. Kisand, L. Haljasmägi, L. Chadli, P. Moingeon, L. Martinez, B. Merrick, K. Bisnauthsing, K. Brooks, M. A. A. Ibrahim, J. Mason, F. Lopez Gomez, K. Babalola, S. Abdul-Jawad, J. Cason, C. Mant, J. Seow, C. Graham, K. J. Doores, F. Di Rosa, J. Edgeworth, M. Shankar-Hari, A. C. Hayday, A dynamic COVID-19 immune signature includes associations with poor prognosis, Nat Med 579, 270–33 (2020).

6. Q.-X. Long, B.-Z. Liu, H.-J. Deng, G.-C. Wu, K. Deng, Y.-K. Chen, P. Liao, J.-F. Qiu, Y. Lin, X.-F. Cai, D.-Q. Wang, Y. Hu, J.-H. Ren, N. Tang, Y.-Y. Xu, L.-H. Yu, Z. Mo, F. Gong, X.-L. Zhang, W.-G. Tian, L. Hu, X.-X. Zhang, J.-L. Xiang, H.-X. Du, H.-W. Liu, C.-H. Lang, X.-H. Luo, S.-B. Wu, X.-P. Cui, Z. Zhou, M.-M. Zhu, J. Wang, C.-J. Xue, X.-F. Li, L. Wang, Z.-J. Li, K. Wang, C.-C. Niu, Q.-J. Yang, X.-J. Tang, Y. Zhang, X.-M. Liu, J.-J. Li, D.-C. Zhang, F. Zhang, P. Liu, J. Yuan, Q. Li, J.-L. Hu, J. Chen, A.-L. Huang, Antibody responses to SARS-CoV-2 in patients with COVID-19, Nat Med 26, 845–848 (2020).

7. C. Cervia, J. Nilsson, Y. Zurbuchen, A. Valaperti, J. Schreiner, A. Wolfensberger, M. E. Raeber, S. Adamo, M. Emmenegger, S. Hasler, P. P. Bosshard, E. De Cecco, E. Bächli, A. Rudiger, M. Stüssi-Helbling, L. C. Huber, A. S. Zinkernagel, D. J. Schaer, A. Aguzzi, U. Held, E. Probst-Müller, S. K. Rampini, O. Boyman, Systemic and mucosal antibody secretion specific to SARS-CoV-2 during mild versus severe COVID-19, bioRxiv doi:2020.05.21.108308 (2020).

8. D. Weiskopf, K. S. Schmitz, M. P. Raadsen, A. Grifoni, N. M. A. Okba, H. Endeman, J. P. C. van den Akker, R. Molenkamp, M. P. G. Koopmans, E. C. M. van Gorp, B. L. Haagmans, R. L. de Swart, A. Sette, R. D. de Vries, Phenotype and kinetics of SARS-CoV-2-specific T cells in COVID-19 patients with acute respiratory distress syndrome, Science Immunology 5, eabd2071 (2020).

9. C. Rydyznski Moderbacher, S. I. Ramirez, J. M. Dan, A. Grifoni, K. M. Hastie, D. Weiskopf, S. Belanger, R. K. Abbott, C. Kim, J. Choi, Y. Kato, E. G. Crotty, C. Kim, S. A. Rawlings, J. Mateus, L. P. V. Tse, A. Frazier, R. Baric, B. Peters, J. Greenbaum, E. Ollmann Saphire, D. M. Smith, A. Sette, S. Crotty, Antigen-Specific Adaptive Immunity to SARS-CoV-2 in Acute COVID-19 and Associations with Age and Disease Severity, Cell 183, 996–1012.e19 (2020).

10. A. T. Tan, M. Linster, C. W. Tan, N. Le Bert, W. N. Chia, K. Kunasegaran, Y. Zhuang, C. Y. L. Tham, A. Chia, G. J. Smith, B. Young, S. Kalimuddin, J. G. H. Low, D. Lye, L.-F. Wang, A. Bertoletti, Early induction of SARS-CoV-2 specific T cells associates with rapid viral clearance and mild disease in COVID-19 patients, bioRxiv doi:10.1101/2020.10.15.341958 (2020).

11. J. Zhao, J. Zhao, S. Perlman, T cell responses are required for protection from clinical disease and for virus clearance in severe acute respiratory syndrome coronavirus-infected mice, Journal of Virology 84, 9318–9325 (2010).

12. J. Zhao, J. Zhao, A. K. Mangalam, R. Channappanavar, C. Fett, D. K. Meyerholz, S. Agnihothram, R. S. Baric, C. S. David, S. Perlman, Airway Memory CD4 + T Cells Mediate Protective Immunity against Emerging Respiratory Coronaviruses, Immunity 44, 1379–1391 (2016).

13. E. Lavezzo, E. Franchin, C. Ciavarella, G. Cuomo-Dannenburg, L. Barzon, C. Del Vecchio, L. Rossi, R. Manganelli, A. Loregian, N. Navarin, D. Abate, M. Sciro, S. Merigliano, E. De Canale, M. C. Vanuzzo, V. Besutti, F. Saluzzo, F. Onelia, M. Pacenti, S. G. Parisi, G. Carretta, D. Donato, L. Flor, S. Cocchio, G. Masi, A. Sperduti, L. Cattarino, R. Salvador, M. Nicoletti, F. Caldart, G. Castelli, E. Nieddu, B. Labella, L. Fava, M. Drigo, K. A. M. Gaythorpe, Imperial College COVID-19 Response Team, A. R. Brazzale, S. Toppo, M. Trevisan, V. Baldo, C. A. Donnelly, N. M. Ferguson, I. Dorigatti, A. Crisanti, Suppression of a SARS-CoV-2 outbreak in the Italian municipality of Vo’, Nature 584, 425–429 (2020).

14. Q.-X. Long, X.-J. Tang, Q.-L. Shi, Q. Li, H.-J. Deng, J. Yuan, J.-L. Hu, W. Xu, Y. Zhang, F.-J. Lv, K. Su, F. Zhang, J. Gong, B. Wu, X.-M. Liu, J.-J. Li, J.-F. Qiu, J. Chen, A.-L. Huang, Clinical and immunological assessment of asymptomatic SARS-CoV-2 infections, Nat Med 63, 706–12 (2020).

15. T. Sekine, A. Perez-Potti, O. Rivera-Ballesteros, K. Strålin, J.-B. Gorin, A. Olsson, S. Llewellyn-Lacey, H. Kamal, G. Bogdanovic, S. Muschiol, D. J. Wullimann, T. Kammann, J. Emgård, T. Parrot, E. Folkesson, Karolinska COVID-19 Study Group, O. Rooyackers, L. I. Eriksson, J.-I. Henter, A. Sönnerborg, T. Allander, J. Albert, M. Nielsen, J. Klingström, S. Gredmark-Russ, N. K. Björkström, J. K. Sandberg, D. A. Price, H. G. Ljunggren, S. Aleman, M. Buggert, Robust T Cell Immunity in Convalescent Individuals with Asymptomatic or Mild COVID-19, Cell 183, 158–168.e14 (2020).

16. C. J. Reynolds, L. Swadling, J. M. Gibbons, C. Pade, M. Jensen, M. O. Diniz, N. M. Schmidt, D. K. Butler, O. E. Amin, S. N. L. Bailey, S. Talyor, J. Jones, M. Jones, W. Y. J. Lee, J. Rosenheim, A. Chandran, G. Joy, C. Di Genova, N. J. Temperton, J. Lambourne, T. Cutino-Moguel, M. Andiapen, M. Fontana, A. Smit, A. Semper, B. O’Brien, B. Chain, T. Brooks, C. Manisty, T. Treibel, J. Moon, COVIDsortium Investigators, M. C. Noursadeghi, COVIDsortium Immune correlates network, D. M. Altmann, M. K. Mani, A. McKnight, R. J. Boyton, Healthcare workers with mild / asymptomatic SARS-CoV-2 infection show T cell responses and neutralising antibodies after the first wave, medRxiv doi:0.1101/2020.10.13.20211763 (2020).

17. X.-N. Zhao, Y. You, G.-L. Wang, H.-X. Gao, X.-M. Cui, L.-J. Duan, S.-B. Zhang, Y.-L. Wang, Lin-Yao. L. Li, J.-H. Lu, H.-B. Wang, J.-F. Fan, H.-W. Zheng, E.-H. Dai, L.-Y. Tian, M.-J. Ma, Longitudinal single-cell immune profiling revealed distinct innate immune response in asymptomatic COVID-19 patients, bioRxiv doi:10.1101/2020.09.02.276865 (2020).

18. C. W. Tan, W. N. Chia, X. Qin, P. Liu, M. I.-C. Chen, C. Tiu, Z. Hu, V. C.-W. Chen, B. E. Young, W. R. Sia, Y.-J. Tan, R. Foo, Y. Yi, D. C. Lye, D. E. Anderson, L.-F. Wang, A SARS-CoV-2 surrogate virus neutralization test based on antibody-mediated blockage of ACE2-spike protein-protein interaction, Nat Biotechnol 395, 470–6 (2020).

19. N. Le Bert, A. T. Tan, K. Kunasegaran, C. Y. L. Tham, M. Hafezi, A. Chia, M. H. Y. Chng, M. Lin, N. Tan, M. Linster, W. N. Chia, M. I.-C. Chen, L.-F. Wang, E. E. Ooi, S. Kalimuddin, P. A. Tambyah, J. G.-H. Low, Y.-J. Tan, A. Bertoletti, SARS-CoV-2-specific T cell immunity in cases of COVID-19 and SARS, and uninfected controls, Nature 584, 457–462 (2020).

20. E. Becht, L. McInnes, J. Healy, C.-A. Dutertre, I. W. H. Kwok, L. G. Ng, F. Ginhoux, E. W. Newell, Dimensionality reduction for visualizing single-cell data using UMAP, Nat Biotechnol 37, 38–44 (2018).

21. C. Atyeo, S. Fischinger, T. Zohar, M. D. Slein, J. Burke, C. Loos, D. J. McCulloch, K. L. Newman, C. Wolf, J. Yu, K. Shuey, J. Feldman, B. M. Hauser, T. Caradonna, A. G. Schmidt, T. J. Suscovich, C. Linde, Y. Cai, D. Barouch, E. T. Ryan, R. C. Charles, D. Lauffenburger, H. Chu, G. Alter, Distinct Early Serological Signatures Track with SARS-CoV-2 Survival, Immunity 53, 524–532.e4 (2020).

22. L. Henss, T. Scholz, C. von Rhein, I. Wieters, F. Borgans, F. J. Eberhardt, K. Zacharowski, S. Ciesek, G. Rohde, M. Vehreschild, C. Stephan, T. Wolf, H. Hofmann-Winkler, H. Scheiblauer, B. S. Schnierle, Analysis of humoral immune responses in SARS-CoV-2 infected patients, The Journal of Infectious Diseases doi:10.1093/infdis/jiaa680 (2020).

23. M. C. Woodruff, R. P. Ramonell, D. C. Nguyen, K. S. Cashman, A. S. Saini, N. S. Haddad, A. M. Ley, S. Kyu, J. C. Howell, T. Ozturk, S. Lee, N. Suryadevara, J. B. Case, R. Bugrovsky, W. Chen, J. Estrada, A. Morrison-Porter, A. Derrico, F. A. Anam, M. Sharma, H. M. Wu, S. N. Le, S. A. Jenks, C. M. Tipton, B. Staitieh, J. L. Daiss, E. Ghosn, M. S. Diamond, R. H. Carnahan, J. E. Crowe, W. T. Hu, F. E.-H. Lee, I. Sanz, Extrafollicular B cell responses correlate with neutralizing antibodies and morbidity in COVID-19, Nat Immunol 21, 1506–16 (2020).

24. J. Zuo, A. Dowell, H. Pearce, K. Verma, H. M. Long, J. Begum, F. Aiano, Z. Amin-Chowdhury, B. Hallis, L. Stapley, R. Borrow, E. Linley, S. Ahmad, B. Parker, A. Horsley, G. Amirthalingam, K. Brown, M. E. Ramsay, S. Ladhani, P. Moss, Robust SARS-CoV-2-specific T-cell immunity is maintained at 6 months following primary infection, bioRxiv doi:0.1101/2020.11.01.362319 (2020).

25. Y. Peng, A. J. Mentzer, G. Liu, X. Yao, Z. Yin, D. Dong, W. Dejnirattisai, T. Rostron, P. Supasa, C. Liu, C. Lopez-Camacho, J. Slon-Campos, Y. Zhao, D. I. Stuart, G. C. Paesen, J. M. Grimes, A. A. Antson, O. W. Bayfield, D. E. D. P. Hawkins, D.-S. Ker, B. Wang, L. Turtle, K. Subramaniam, P. Thomson, P. Zhang, C. Dold, J. Ratcliff, P. Simmonds, T. de Silva, P. Sopp, D. Wellington, U. Rajapaksa, Y.-L. Chen, M. Salio, G. Napolitani, W. Paes, P. Borrow, B. M. Kessler, J. W. Fry, N. F. Schwabe, M. G. Semple, J. K. Baillie, S. C. Moore, P. J. M. Openshaw, M. A. Ansari, S. Dunachie, E. Barnes, J. Frater, G. Kerr, P. Goulder, T. Lockett, R. Levin, Y. Zhang, R. Jing, L.-P. Ho, Oxford Immunology Network Covid-19 Response T cell Consortium, ISARIC4C Investigators, R. J. Cornall, C. P. Conlon, P. Klenerman, G. R. Screaton, J. Mongkolsapaya, A. McMichael, J. C. Knight, G. Ogg, T. Dong, Broad and strong memory CD4+ and CD8+ T cells induced by SARS-CoV-2 in UK convalescent individuals following COVID-19, Nat Immunol 21, 1336–1345 (2020).

26. Y. Wang, L. Zhang, L. Sang, F. Ye, S. Ruan, B. Zhong, T. Song, A. N. Alshukairi, R. Chen, Z. Zhang, M. Gan, A. Zhu, Y. Huang, L. Luo, C. K. P. Mok, M. M. Al Gethamy, H. Tan, Z. Li, X. Huang, F. Li, J. Sun, Y. Zhang, L. Wen, Y. Li, Z. Chen, Z. Zhuang, J. Zhuo, C. Chen, L. Kuang, J. Wang, H. Lv, Y. Jiang, M. Li, Y. Lin, Y. Deng, L. Tang, J. Liang, J. Huang, S. Perlman, N. Zhong, J. Zhao, J. S. Malik Peiris, Y. Li, J. Zhao, Kinetics of viral load and antibody response in relation to COVID-19 severity, J. Clin. Invest. 130, 5235–5244 (2020).

27. C. Gaebler, Z. Wang, J. C. C. Lorenzi, F. Muecksch, S. Finkin, M. Tokuyama, M. Ladinsky, A. Cho, M. Jankovic, D. Schaefer-Babajew, T. Y. Oliveira, M. Cipolla, C. Viant, C. O. Barnes, A. Hurley, M. Turroja, K. Gordon, K. G. Millard, V. Ramos, F. Schmidt, Y. Weisblum, D. Jha, M. Tankelevich, J. Yee, I. Shimeliovich, D. F. Robbiani, Z. Zhao, A. Gazumyan, T. Hatziioannou, P. J. Bjorkman, S. Mehandru, P. D. Bieniasz, M. Caskey, M. C. Nussenzweig, Evolution of Antibody Immunity to SARS-CoV-2, bioRxiv doi:10.1101/2020.11.03.367391 (2020).

28. T. Ford, C. Wenden, A. Mbekeani, L. Dally, J. H. Cox, M. Morin, N. Winstone, A. V. S. Hill, J. Gilmour, K. J. Ewer, Cryopreservation-related loss of antigen-specific IFNγ producing CD4+ T-cells can skew immunogenicity data in vaccine trials: Lessons from a malaria vaccine trial substudy, Vaccine 35, 1898–1906 (2017).

29. A. Grifoni, D. Weiskopf, S. I. Ramirez, J. Mateus, J. M. Dan, C. R. Moderbacher, S. A. Rawlings, A. Sutherland, L. Premkumar, R. S. Jadi, D. Marrama, A. M. de Silva, A. Frazier, A. F. Carlin, J. A. Greenbaum, B. Peters, F. Krammer, D. M. Smith, S. Crotty, A. Sette, Targets of T Cell Responses to SARS-CoV-2 Coronavirus in Humans with COVID-19 Disease and Unexposed Individuals, Cell 181, 1489–1501.e15 (2020).

30. J. S. Lee, S. Park, H. W. Jeong, J. Y. Ahn, S. J. Choi, H. Lee, B. Choi, S. K. Nam, M. Sa, J.-S. Kwon, S. J. Jeong, H. K. Lee, S. H. Park, S.-H. Park, J. Y. Choi, S.-H. Kim, I. Jung, E.-C. Shin, Immunophenotyping of COVID-19 and influenza highlights the role of type I interferons in development of severe COVID-19, Science Immunology 5, eabd1554 (2020).

31. C. Lucas, P. Wong, J. Klein, T. B. R. Castro, J. Silva, M. Sundaram, M. K. Ellingson, T. Mao, J. E. Oh, B. Israelow, T. Takahashi, M. Tokuyama, P. Lu, A. Venkataraman, A. Park, S. Mohanty, H. Wang, A. L. Wyllie, C. B. F. Vogels, R. Earnest, S. Lapidus, I. M. Ott, A. J. Moore, M. C. Muenker, J. B. Fournier, M. Campbell, C. D. Odio, A. Casanovas-Massana, Yale IMPACT Team, R. Herbst, A. C. Shaw, R. Medzhitov, W. L. Schulz, N. D. Grubaugh, C. Dela Cruz, S. Farhadian, A. I. Ko, S. B. Omer, A. Iwasaki, Longitudinal analyses reveal immunological misfiring in severe COVID-19, Nature 584, 463–469 (2020).

32. S. De Biasi, M. Meschiari, L. Gibellini, C. Bellinazzi, R. Borella, L. Fidanza, L. Gozzi, A. Iannone, D. Lo Tartaro, M. Mattioli, A. Paolini, M. Menozzi, J. Milić, G. Franceschi, R. Fantini, R. Tonelli, M. Sita, M. Sarti, T. Trenti, L. Brugioni, L. Cicchetti, F. Facchinetti, A. Pietrangelo, E. Clini, M. Girardis, G. Guaraldi, C. Mussini, A. Cossarizza, Marked T cell activation, senescence, exhaustion and skewing towards TH17 in patients with COVID-19 pneumonia, Nature Communications 11, 1199–17 (2020).

33. M.-S. Rha, H. W. Jeong, J.-H. Ko, S. J. Choi, I.-H. Seo, J. S. Lee, M. Sa, A. R. Kim, E.-J. Joo, J. Y. Ahn, J. H. Kim, K.-H. Song, E. S. Kim, H. B. Kim, Y. K. Kim, S.-H. Park, J. Y. Choi, K. R. Peck, E.-C. Shin, IFN-γ is Produced by Pd-1 + Cells Among SARS-CoV-2-Specific MHC-I Multimer+CD8 + T Cells in Acute and Convalescent COVID-19 Patients. Available at SSRN: https://ssrn.com/abstract=3684758 (2020).

34. K. N. Couper, D. G. Blount, E. M. Riley, IL-10: the master regulator of immunity to infection, J. Immunol. 180, 5771–5777 (2008).

35. J. Sun, R. Madan, C. L. Karp, T. J. Braciale, Effector T cells control lung inflammation during acute influenza virus infection by producing IL-10, Nat Med 15, 277–284 (2009).

36. A. Grifoni, J. Sidney, Y. Zhang, R. H. Scheuermann, B. Peters, A. Sette, A Sequence Homology and Bioinformatic Approach Can Predict Candidate Targets for Immune Responses to SARS-CoV-2, Cell Host and Microbe 27, 671–680.e2 (2020).

